# Cortical and subcortical brain networks predict prevailing heart rate

**DOI:** 10.1101/2023.09.23.559114

**Authors:** Amy Isabella Sentis, Javier Rasero, Peter J. Gianaros, Timothy D. Verstynen

**Author notes:** **Corresponding author:** Timothy Verstynen, 342E Baker Hall Department of Psychology Carnegie Mellon University 5000 Forbes Avenue, Pittsburgh, PA 15213. Co-senior authors (Peter J. Gianaros, Timothy D. Verstynen).

## Abstract

Resting heart rate may confer risk for cardiovascular disease (CVD) and other adverse cardiovascular events. While the brainstem’s autonomic control over heart rate is well established, less is known about the regulatory role of higher-level cortical and subcortical brain regions, especially in humans. The present study sought to characterize the brain networks that predict variation in prevailing heart rate in otherwise healthy adults. We used machine learning approaches designed for complex, high-dimensional datasets, to predict variation in instantaneous heart period (the inter-heartbeat-interval) from whole brain hemodynamic signals measured by fMRI. Task-based and resting-state fMRI, as well as peripheral physiological recordings, were taken from two datasets that included extensive repeated measurements within individuals. Our models reliably predicted instantaneous heart period from whole brain fMRI data both within and across individuals, with prediction accuracies being highest when measured within-participants. We found that a network of cortical and subcortical brain regions, many linked to psychological stress, were reliable predictors of variation in heart period. This adds to evidence on brain-heart interactions and constitutes an incremental step towards developing clinically-applicable biomarkers of brain contributions to CVD risk.

**Impact statement:** Using whole brain fMRI data, we reliably predicted instantaneous heart period within and across individuals from the activity of a network of cortical and subcortical brain regions, many linked to psychological stress. This adds to existing evidence on brain-heart interactions and constitutes a step towards developing clinically-applicable biomarkers of brain contributions to CVD risk.

## 1. Introduction

Resting heart rate (HR) is not only a predictor of all-cause mortality, but is also a risk factor for cardiovascular disease in individuals with and without pre-existing cardiovascular disease (CVD; [1–4]). High resting HR has been associated with progression of coronary artery atherosclerosis and the occurrence of myocardial ischemia and arrhythmias, and is implicated in left ventricular dysfunction [2–7]. Conversely, reduction of resting HR has long been associated with the prevention of activity related angina and ischemia [3]. High resting HR is often comorbid in individuals with other cardiometabolic risk factors, including hypertension, high blood lipid and glucose levels, and overweight or high BMI [3,8]. A large scale, long-term follow up epidemiological study found high resting HR to be a risk factor for all-cause mortality, as well as death from acute myocardial infarction, after adjusting for possible confounds such as age, BMI, systolic BP, diabetes diagnosis and level of physical activity [3,9].

Evidence from animal studies and human lesion studies have shown that HR is under autonomic control, with the nucleus of the solitary tract in the brainstem exerting proximal regulation [10–14]. This brain-heart link has been more recently validated by human neuroimaging studies [12,15]. This link also extends further “up” the brain, to evolutionarily newer brain regions like the telencephelon (i.e., neocortex), which seems to play a role in stress-related modulation of HR [16]. During psychologically stressful contexts, the cortical areas appear to exert direct and indirect cardiac control via the autonomic nervous system (specifically, sympathetic activation followed by parasympathetic inhibition) [17–20]. Indeed, the role of psychological stress in modulating cardiac function provides a particularly clear indication of the role that cortical function has on HR [16,21–24].

Here we sought to characterize the brain networks that predict variation in prevailing HR within healthy individuals. To do so, we used machine learning approaches designed for complex and high-dimensional datasets to predict instantaneous heart period (the inter-heartbeat-interval) from temporal variation in whole brain hemodynamic signals measured using fMRI. Here we evaluated two hypotheses. First, we evaluated whether it is possible to reliably predict modulation of heart period from hemodynamic responses in the brain *within individuals*. Second, we looked at whether the brain regions that are most important for this prediction would encompass part or all of visceral control circuits.

## 2. Methods

### 2.1 Human QA dataset

#### 2.1.1 Participants

Neuroimaging and cardiovascular data were collected from one healthy participant (white male in his forties) at 14 repeated scan sessions over a period of 20 weeks at the CMU-Pitt BRIDGE Center (RRID:SCR_023356). Three fMRI scans with concurrent electrocardiogram (ECG) and pneumatic belt physiological signals were collected at each session, along with a structural MRI scan, for a total of 42 runs. The participant provided informed consent to complete the study, which was approved by the Carnegie Mellon University (Pittsburgh, PA) Institutional Review Board.

#### 2.1.2 MRI data acquisition and processing

##### Acquisition

Functional blood oxygenation level-dependent images were collected on a Siemens Prisma 3 Tesla scanner, equipped with a 64-channel head coil. Over a 8 minute 54 second period with eyes open, resting-state and task dependent functional images were acquired with acquisition parameters as follows: 2mm iso voxels, FOV = 212x212mm, TR = 1500ms, TE = 30ms, FA = 79°, multiband acceleration factor = 4. Sixty-eight interleaved slices (2mm thickness, no gap) in the ascending direction were obtained for each of 353 volumes (with three initial volumes discarded to allow for magnetic equilibration). T1-weighted neuroanatomical magnetization prepared rapid gradient echo (MPRAGE) images were collected in 208 slices (1mm thickness, no gap) over a 7 minute 58 second period with acquisition parameters as follows: 1mm iso voxels, FOV = 256x256mm, TR = 2300ms, TE = 2.03ms, TI = 900ms, FA = 9°.

##### Preprocessing

Preprocessing was performed using fMRIPrep version 20.2.7 ([25]; RRID:SCR_016216). Eight T1-weighted images collected across the 14 sessions were combined into one average T1-weighted reference map. Anatomical and functional images were normalized to the ICBM 152 Nonlinear Asymmetrical template version 2009c ([26], RRID:SCR_008796; TemplateFlow ID: MNI152NLin2009cAsym) to facilitate between-participant and between-dataset comparisons.

##### Gray Matter (GM) mask

Using the GM tissue probability reference image in MNI152NLin2009cAsym standard space output from fMRIPrep, we generated a binary mask (thresholded at probability > 0.2) to limit our analysis to GM voxels.

#### 2.1.3 Heart period from ECG

ECG data were collected concurrently during functional MRI (fMRI) scans at a sampling rate of 400 Hz using the Siemens physiological monitoring unit’s standard configuration. Data were processed using *niphlem* (https://coaxlab.github.io/niphlem/) and included the following steps: 1) subtraction of ground ECG channel from remaining channels, 2) demean channels, 3) bandpass filtering (0.6 Hz - 5 Hz), 4) average channels. Signal peaks were identified (specifically, the R peaks within the QRS waveform) with subsequent detection of artifacts through two one-sided Grubb’s tests for outliers and correction. The processed ECG signal peaks were then downsampled to match the fMRI TR, to allow for a one-to-one correspondence of our X and y data for our prediction models.

It is known that heart rate behaves non-linearly in many contexts, especially with respect to underlying autonomic input [27,28]. An alternative measure is heart period (the interbeat interval, or the time between ECG signal peaks), which is the inverse of heart rate. While both measures are controlled in part by autonomic inputs, there exists a more linear relationship between autonomic control and heart period, when compared to heart rate [27,28]. Thus heart rate and heart period are not interchangeable with respect to autonomic control. For this reason, we used heart period as our target variable.

#### 2.1.4 Analysis

Figure 1 shows the structure of our analysis pipeline, which involves three stages: preprocessing, denoising, and model analysis. Preprocessing was performed using fMRIPrep and *niphlem* for the fMRI and physiological data, respectively as detailed in Sections 2.1.2 and 2.1.3. Code is available at https://github.com/CoAxLab/dynamic-hr.

**Figure 1:**
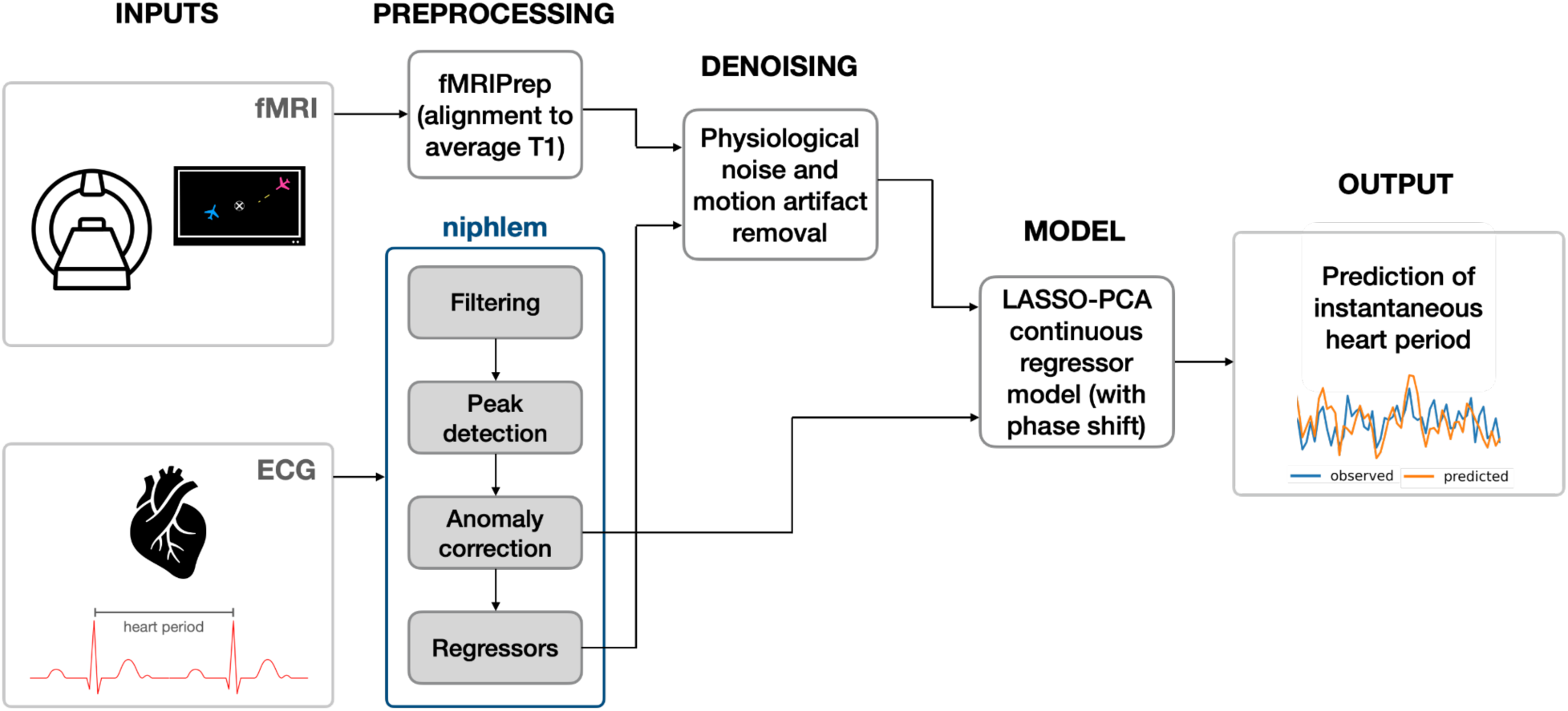
Analysis pipeline schematic.

Denoising constituted removal of physiological noise and motion artifacts (rotational and translational) from the hemodynamic signal itself. To do so, we used *niphlem* to generate variability regressors from the cardiac and respiratory response functions [29,30]. We then conducted a GLM of these variability regressors onto our fMRI signal to capture the artifact components from mechanical physiological noise. We removed these artifacts from the fMRI signal by isolating the GLM residuals to use for further analysis.

The next stage of our pipeline was a leave-one-run-out nested cross-validated LASSO-PCA model that took the GLM residuals as input and predicted instantaneous heart period. We set up multiple models (13 total), each of which predicted instantaneous heart period from fMRI timeseries data at time points ranging from t-3 seconds to t+15 seconds (shifting by one TR at a time, i.e., 1.5 seconds). This range of lag time shifts between the fMRI signal and instantaneous heart period ensures that we take into account the HRF delay (6-8 seconds) inherent with BOLD data. See Figure 2 for a graphical illustration of the lag time shifts to isolate the desired drive signal. Thus, for each run across sessions, instantaneous heart period was predicted at every TR from GM voxels of the fMRI signal. First, as part of the PCA step, singular value decomposition was performed on the training runs to reduce the dimensionality of the predictor matrix. This also addresses the issue of multicollinearity across voxels. The inner loop then performed five-fold cross-validation on the training runs to optimize lambda (𝝀), the shrinkage parameter that controls the L1 penalty in LASSO, using a sequence of 100 𝝀 values. Finally, the outer leave-one-run-out cross validation loop performed the entire LASSO-PCA algorithm using this optimal lambda value. We repeated this analysis using the preprocessed timeseries before removal of artifacts for comparison.

**Figure 2:**
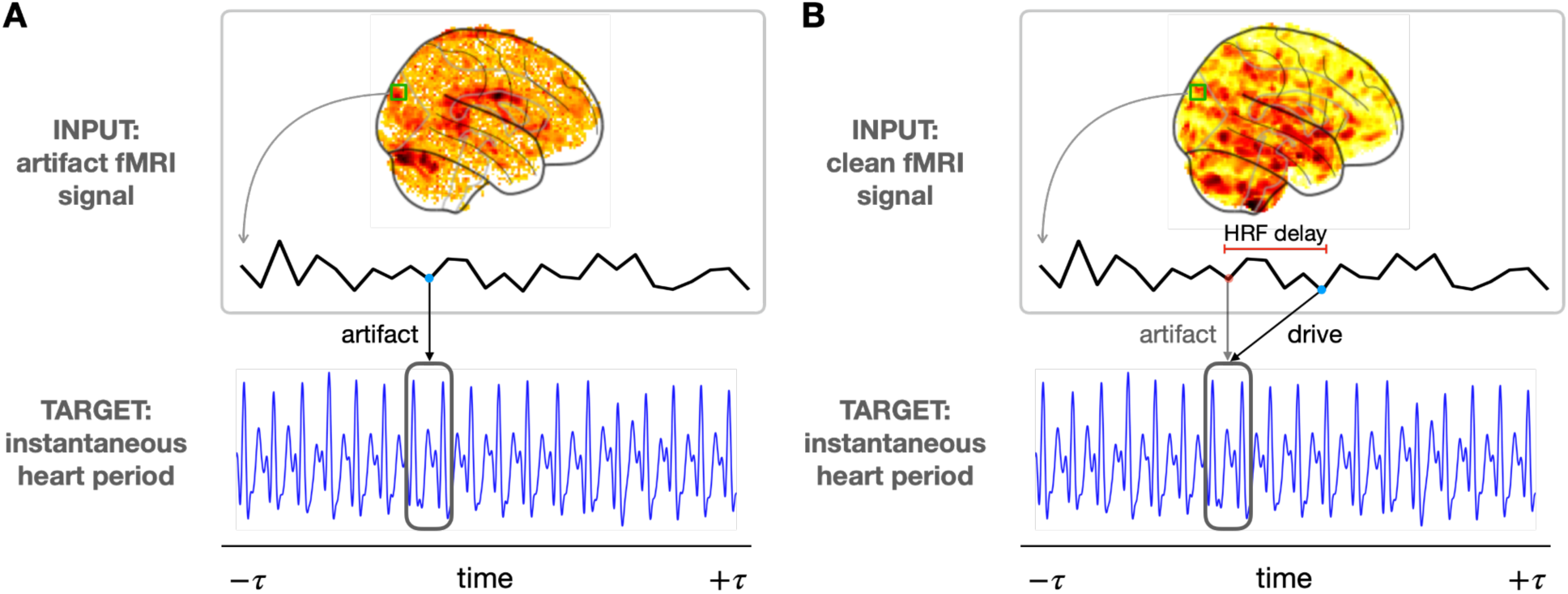
A) Prediction of heart period from physiological noise component of fMRI signal. B) Prediction of heart period after removing artifacts from the fMRI signal and accounting for the HRF delay.

Overall model performance was evaluated using prediction accuracy on a hold-out test set. This was done by comparing predicted heart period and observed heart period using fMRI runs that were not included in the model training. Model performance was measured using the Pearson correlation coefficient, r, between the heart period predicted from the brain responses and the observed heart period.

In order to visualize which brain regions were contributing the most to the prediction of heart period, we projected the voxelwise decoding maps into approximations of their encoding representations [31]. For this LASSO coefficients were extracted from each model run and multiplied by the V matrix (from the singular value decomposition 𝑋 = 𝑈𝑆𝑉 ^𝑇^) to generate weights in feature space. These weights were then multiplied by the covariance matrix of X (our voxel data) to convert to encoding weights [31].

To minimize computation time and excessive memory requirements of our analysis pipeline with large datasets, we adopted a modular approach that takes advantage of model averaging [32].

We trained individual LASSO-PCA models for each run and extracted the associated weights. For the outer leave-one-out cross-validation loop, we averaged these weights from the n-1 trained models, using the mean voxel decoding weights, to test our model’s hold-out prediction accuracy.

We performed a two-sided one-sample t-test on the encoding weights for lag time shift +7.5 seconds for each run with a false discovery rate (FDR) < 0.00005 to generate a statistically thresholded weight map corresponding to the Human QA dataset participant. This specific lag was chosen because it represents the optimal timepoint for detecting the blood oxygen level dependent (BOLD) response evoked from underlying neural activity, given the sampling rate of the data set. Visualization of the participant’s projected weights on the 3D brain was performed using Surf Ice (https://www.nitrc.org/projects/surfice/).

#### 2.1.5 Power analysis

We conducted a power analysis to test the number of runs needed for reliable prediction of heart period from fMRI data, using sample sizes n = 2, 4, 8, 10, 16, 20, and 30. For each sample size, we randomly selected n runs from the 42 total runs to use in our analysis. Working with the cleaned fMRI data at lag time shift +7.5 seconds, we trained a LASSO-PCA model on each run then used a leave-one-out cross-validation scheme to test each model on the average weights from the n-1 trained models, as described in Section 2.1.4. This procedure was repeated 40 times for each sample size. Hold-out test set prediction accuracy was evaluated using Pearson correlation coefficient and the mean r value and 95% confidence interval was calculated for each sample size.

### 2.2 Natural Scenes Dataset (NSD)

#### 2.2.1 Participants

Neuroimaging and cardiovascular data from eight healthy participants (six females, two males; aged 19–32 years; three Asian, five White) from the NSD were used for Experiment 2. A detailed description of the Natural Scenes Dataset (NSD; http://naturalscenesdataset.org) is provided elsewhere [33]. Physiological data was only available for a subset of scan sessions: four participants had data from ten scan sessions (S1, S2, S5, S7; nsd21-30), two participants had data from nine scan sessions (S3, S6; nsd21-29) and two participants had data from seven scan sessions (S4, S8; nsd21-27). Each session contained 14 functional runs (12 task and two resting-state scans). The total number of runs for each participant are as follows: S1 n = 140, S2 n = 140, S3 n = 126, S4 n = 93, S5 n = 140, S6 n = 126, S7 n = 139, and S8 n = 84.

#### 2.2.2 Heart period from pulse-oximetry

Physiological data from pulse-oximetry and pneumatic belt were used to record cardiac and respiration events respectively. *Niphlem* was used as described in Section 2.1.2 to extract instantaneous heart period and generate cardiac and respiratory variability regressors. In this case however, heart period was derived from the pulse-oximetry data.

Runs with noisy or interrupted pulse-oximetry data that prevented recovery of heart period were excluded from analysis: five runs for S4, one run for S7, and 14 runs (one entire session) for S8.

#### 2.2.3 MRI data acquisition and processing

##### Acquisition

FMRI data in the NSD were collected at 7T using a whole-brain, 1.8mm, 1.6 second, gradient-echo, echo-planar imaging (EPI) pulse sequence. Further details can be found in [33].

##### Preprocessing

We used the 1.8mm volume preparation of the preprocessed NSD timeseries data. FMRI preprocessing involved two steps. First, a temporal resampling was performed using a cubic interpolation. The timeseries for each voxel was upsampled to 1.333 second to correct for slice-time differences (resulting in 226 volumes for each run). Second, a spatial resampling was performed using a cubic interpolation to correct for head motion, EPI distortion, gradient nonlinearities, and across-scan-session alignment.

##### GM mask

We used the surface-based HCP_MMP1 parcellation [34] available in 1.8mm functional space for each participant to generate binary masks that limited our analyses to GM voxels.

#### 2.2.4 Individual participant analysis

Preprocessing, denoising and model analyses for the NSD data were performed separately for each participant using the methods detailed in Section 2.1.4. We again set up 13 models in total, each of which predicted instantaneous heart period from fMRI timeseries data at time points ranging from t-2.66 seconds to t+13.3 seconds (shifting by one TR at a time, i.e., 1.33 seconds).

We performed one-sampled, two-sided t-tests on the encoding weights for lag time shift +7.99 seconds across runs for each participant with FDR < 0.05. For three participants we were able to use more conservative thresholds: S3=0.0001, S5=0.005, and S7=0.001.

#### 2.2.5 Group analysis

We tested each participant’s average trained model on every other participant for lag time shift +7.99 seconds. In order to do so, we first converted each participant’s GM mask into MNI space (using the nsdcode mapping utility; https://pypi.org/project/nsdcode/) and then created a global participant mask from the union of the individual masks in MNI space. After converting the individual participant weight maps back into nifti images, we used the union MNI GM mask to extract matrices of a common size across participants. These weight matrices were averaged for each participant. Each average weight matrix was used as the trained model and tested on each individual run for every other participant, resulting in 56 group models. For each of these group models, we calculated the Euclidean distance between weight maps as a measure of similarity. We also replicated the within participant analysis in MNI space for lag time shift +7.99 seconds.

To visualize the common brain regions involved in prediction of heart period across participants, we generated two probability maps (one for positive weights and one for negative weights) that show the probability of a voxel being significant across participants. To do so, we thresholded and then binarized the positive and negative weights from the individual participant t-tests separately, using FDR < 0.05. We then averaged the positive and negative weight maps separately across participants. Each voxel has a value between 0 and 1 that represents the probability of that voxel being significant across all participants. Positive and negative probability maps were overlaid and visualized using Surf Ice.

### 2.3 Generalization

We conducted a cross-dataset analysis to test the generalizability of our method and models, most notably across MRI acquisition resolutions. Using the NSD union MNI GM mask described in Section 2.2.5, we repeated the physiological denoising steps discussed in Section 2.1.4 on the Human QA dataset to extract the GLM residuals (X data) using an identical voxel mapping for both datasets. We then averaged the model decoding weights from lag time shift +7.99 seconds for all runs across all NSD participants to use as our average trained model. Finally, we tested each Human QA dataset run at lag time shift +7.5 seconds on the average NSD trained model, resulting in 42 total models. Model accuracy was again evaluated using Pearson correlation coefficient, along with mean and confidence interval summary measures.

## 3. Results

### 3.1 Within-participant decoding performance

#### 3.1.1 Prediction of instantaneous heart period

Results for the single-participant dataset from our leave-one-run-out cross-validation pipeline can be seen in Figure 3. Here we show our LASSO-PCA model prediction accuracy in the form of the correlation between observed and predicted heart period using clean, or denoised data (i.e. the residuals from the GLM of cardiac and respiration variability regressors), and the raw data (i.e., without the artifact signal from the physiological noise removed). When we look at how well our model predicts instantaneous heart period across lag time shifts, (Figure 3A), we see several patterns emerge. First, the cleaned data (blue lines), with the physiological noise artifacts removed, performs generally much better across lag time shifts than the raw data (orange lines), without the physiological artifacts removed (but otherwise identical). Second, we see the expected peak in prediction accuracy around the 0 and +1.5 second lag time shift, which reflects the point at which the physiological artifacts from cardiac and respiratory events would be expressed in the hemodynamic signal (see Figure 2A). Finally, there is an expected peak in model performance for the cleaned data, but not for the raw data, at lag time shift +7.5 seconds, as represented by the dashed vertical line in panel A, which accounts for any potential drive signals in the brain that regulate variation in instantaneous heart period. In particular, we only see this boost in model performance at this time shifted lag in the data where physiological artifacts have been removed from the signal. Figure 3B shows the instantaneous heart period timeseries from a single run in green, with the predicted heart period from the cleaned data model at lag time shift +7.5 seconds overlaid in purple. The close tracking of the predicted heart period timeseries with the observed instantaneous heart period visually demonstrates the peak in model performance associated with the brain’s drive signal which regulates instantaneous heart period.

**Figure 3:**
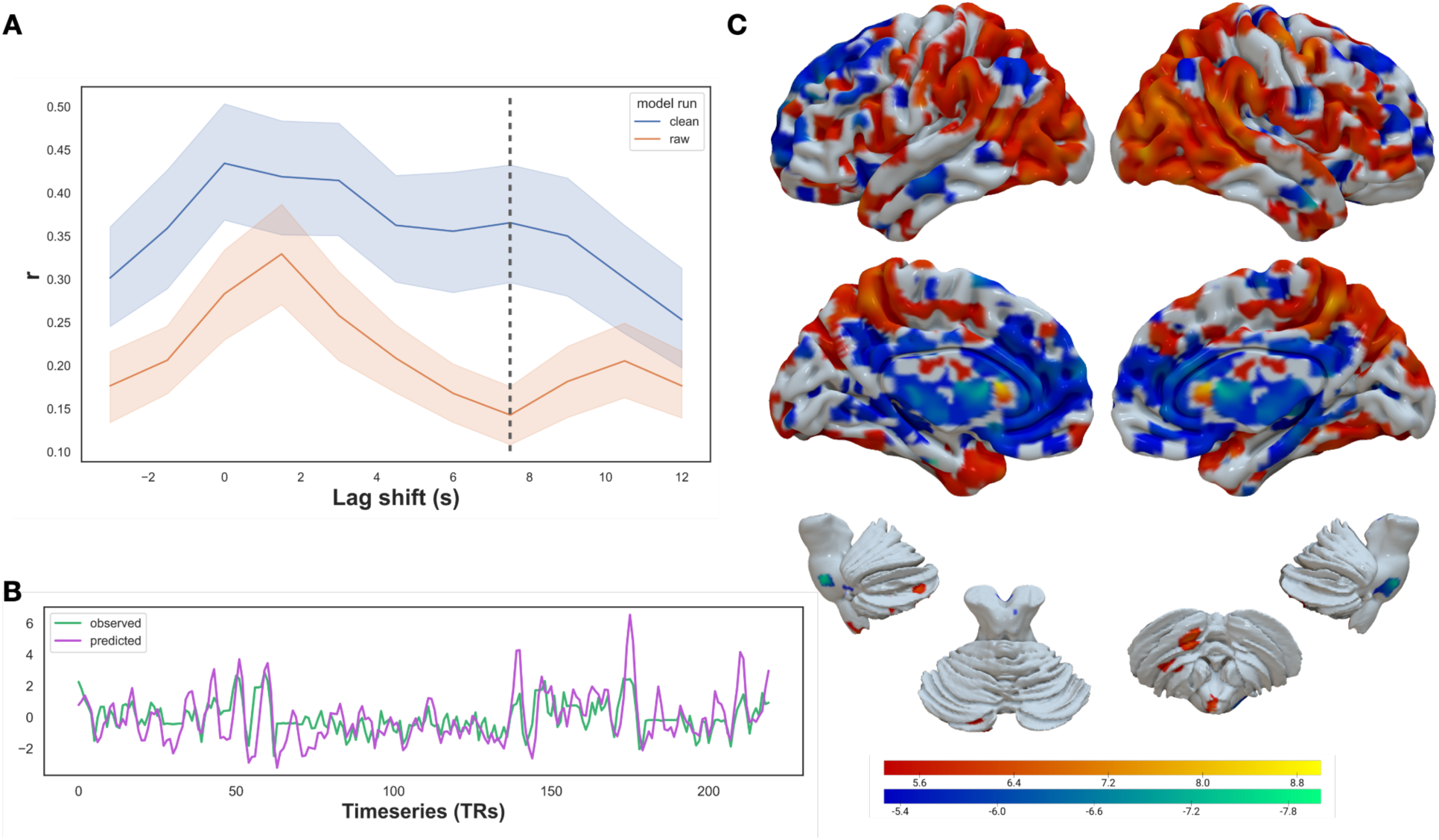
A) Mean out-of-sample Pearson correlation coefficient, r, of predicted vs. observed instantaneous heart period across 14 sessions (42 runs total) for each lag time shift for the original (orange) and cleaned (blue) fMRI signal. Shaded regions represent 95% confidence intervals (calculated using 1000 bootstrap iterations). Dashed vertical line represents the drive signal at lag time shift +7.5 seconds. B) Example observed and predicted instantaneous heart period across the timeseries at time shift +7.5 seconds for a representative run. C) One-sample t-test (FDR correction < 0.00005) of encoding weight maps of instantaneous heart period prediction across sessions and runs for the Human QA dataset participant at time shift +7.5 seconds. Positive weights are shown in red-yellow. Negative weights are shown in blue-green.

These results rely on a large, with-participant dataset to predict instantaneous heart period from variations in the whole brain hemodynamic signal (i.e., 42 fMRI runs collected across 14 separate sessions). One obvious question that arises is: is a dataset this large necessary to predict instantaneous heart period within an individual? In order to determine the number of required runs needed for reliable prediction of heart period, we conducted a power analysis using sample sizes of n = 2, 4, 8, 10, 16, 20, and 30 runs at lag time shift +7.5 seconds. As shown in Supplementary Figure 2, the correlation between observed and predicted heart period increases with increasing sample sizes, plateauing at approximately 16 runs, with a prediction accuracy of r = 0.35. However, the largest jump in prediction accuracy occurs when increasing from sample size of 2 runs to 4 runs, with a greater than 0.1 increase in Pearson correlation coefficient, r. Sample sizes greater than n = 8 show diminishing increases in prediction accuracy. Thus, it seems feasible to develop a reliable predictor of within-participant heart period variation from 1-2 sessions of fMRI scan time, without further optimization of the acquisition protocols (e.g., increased sampling rate, improved shimming).

#### 3.1.3 Areas that associate with instantaneous heart period

Now that we have shown it is possible to reliably predict heart period from whole brain hemodynamic responses, we would like to know which areas are contributing to this prediction. Figure 3C shows the encoding maps, derived from the decoding weights, that reflect the brain regions that positively (red-yellow) and negatively (blue-green) associate with downstream changes in instantaneous heart period. Brain regions including the bilateral occipital cortex, superior parietal cortex, temporal pole, precuneus, supramarginal, dorsolateral prefrontal cortex (dlPFC), right insula, left cerebellum and a portion of the left medial PFC (mPFC) were positively associated with heart period prediction. Specifically, for these regions, increases and or decreases in fMRI BOLD signal were associated with corresponding increased and or decreased instantaneous heart period. Bilateral occipital, superior parietal, supramarginal, temporal pole, superior temporal and temporoparietal junction appear to have the strongest positive association with heart period prediction. Comparatively, bilateral superior frontal, ventromedial PFC (vmPFC), middle temporal, anterior cingulate, angular gyrus, thalamus and periaqueductal gray (PAG) regions were negatively associated with prediction of heart period. For these brain regions, there was an anticorrelated relationship between fMRI BOLD signal and instantaneous heart period, e.g., an increase in the BOLD signal corresponded to a decrease in the instantaneous heart period. Bilateral anterior cingulate, middle temporal, left vmPFC and right posterior cingulate seem to have the strongest negative association with heart period prediction.

### 3.2 Across-participant performance

#### 3.2.1 Replicating single-participant results

In order to replicate our within-participant analysis and extend results to characterizing between-participant performance, we re-ran our model on the eight participants that make up the NSD (see Section 2.2.1). For this we repeated the same analysis shown in Section 3.1 for each NSD participant across lag time shifts −2.67 seconds <= t <= 13.3 seconds with the same LASSO-PCA method as used for the Human QA dataset. Figure 4A shows the individual model accuracies for each NSD participant as quantified by the correlation between observed and predicted heart period across lag time shifts. We see an expected peak around lag time shift +1.33 seconds, as indicated by the leftmost dashed vertical line, that reflects the time window during which physiological artifacts manifest in the BOLD signal. Three participants (S1, S6, S7), plotted in different shades of gray, have unexpectedly lower artifact signal correlation values (i.e., poor prediction at lag time shifts that should recover the physiological artifacts themselves). The rightmost dashed vertical line on the right at lag time shift +7.99 seconds represents the drive signal and is closest timepoint to the identified drive signal from the Human QA dataset at lag time shift +7.5 seconds. Focusing on how well the individual participant models predict instantaneous heart rate as part of the drive signal that regulates variation in heart period at lag time shift +7.99 seconds, we observed some variation across participants. The model accuracies for a majority of participants’ fall in the 0.15 <= r <= 0.2 range (S2, S4-8), while S3 has a larger mean correlation value of r = 0.3065 and S1 has the smallest mean correlation value, of r = 0.0609. However, these patterns also persist across lag time shifts. It is also interesting to note that peak model performance associated with the drive signal does not appear to be located at lag time shift +7.99 seconds for most participants, and indeed the peak performance for NSD participants varies somewhat across time shifts. For example, for participants S3 and S8, the peak performance during the time window associated with the drive signal occurs at +5.33 seconds, while for S4 and S7, it occurs at +6.67 seconds. Additionally, participants S3, S4 and S8 have more obvious drive signal peaks than participants.

**Figure 4:**
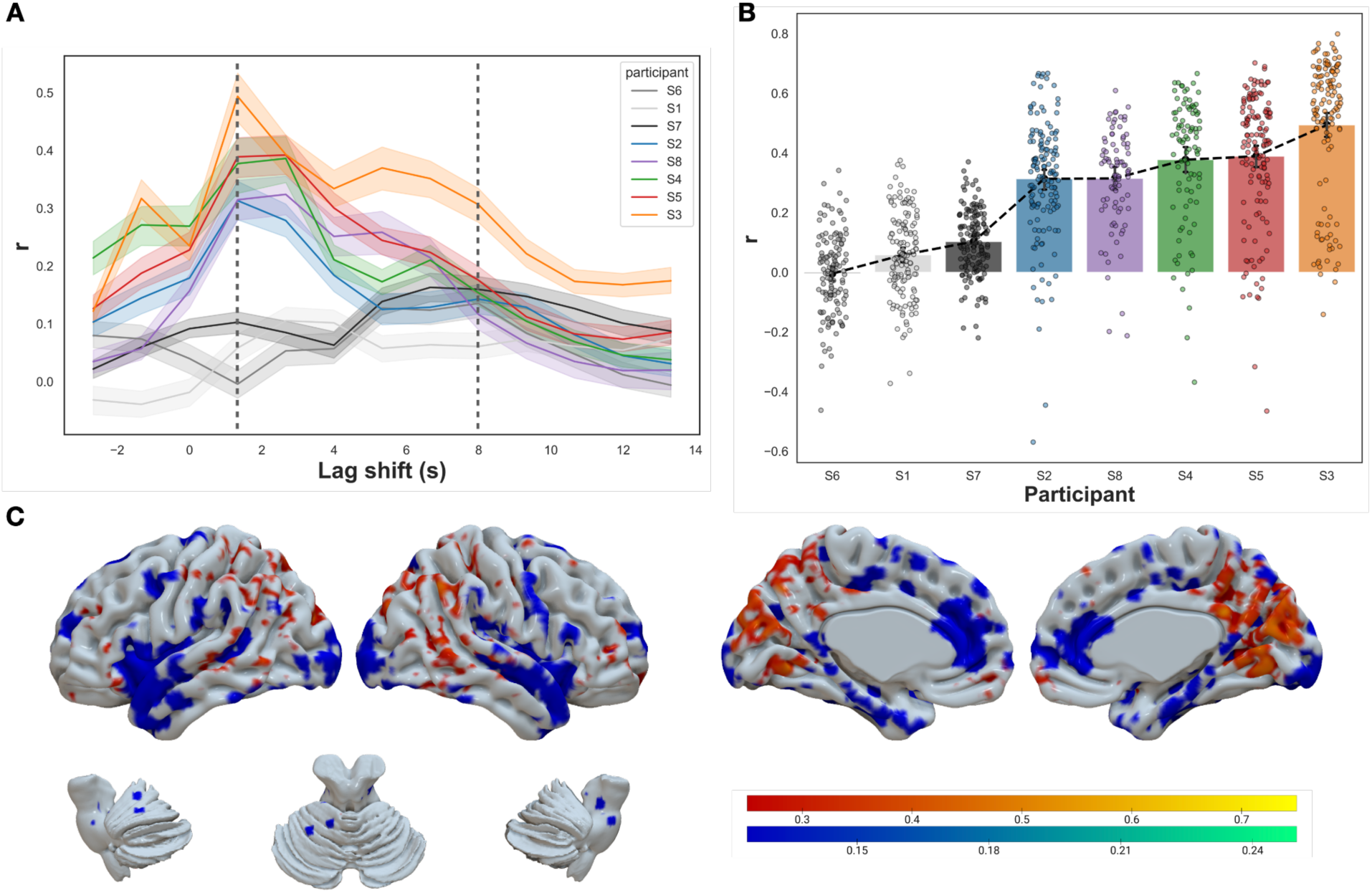
A) Mean out-of-sample Pearson correlation coefficient, r, of instantaneous heart period for each participant for each lag time shift for the clean fMRI signal. Shaded regions represent 95% confidence intervals (calculated using 1000 bootstrap iterations). Dashed vertical lines represent the artifact signal (left) at leg time shift +1.33 seconds and the drive signal (right) at lag time shift +7.99 seconds. B) Correlation coefficients for each run for each participant at lag time shift +1.33 seconds, reflected as the leftmost dashed line in panel A. Bars represent the mean r values, error bars show the 95% confidence intervals. C) Probability map of encoding weights of instantaneous heart period prediction averaged across all participants at lag time shift +7.99 seconds. Positive weights are shown in red-yellow. Negative weights are shown in blue-green.

It is clear from Figure 4A that most, but not all, participants show reliable model performance. However, three participants appear to have overall low prediction accuracy, even in the baseline condition of predicting the time-window when the physiological artifacts are present. Figure 4B shows the prediction accuracy from the window that predicts the artifact signal, at lag time shift +1.33 seconds, as shown by the leftmost dashed vertical line in panel A. For each participant (along the x axis), the correlation between observed and predicted heart period for every run is plotted as a separate point, while the bar shows the mean value and the error bars are 95% confidence intervals from 1000 bootstrap iterations. Participants are plotted in ascending order from left to right: S6 mean r = −0.0044, S1 mean r = 0.0582, S7 mean r = 0.1028, S2 mean r = 0.3133, S8 mean r = 0.3147, S4 mean r = 0.3773, S5 mean r = 0.389, S3 mean r = 0.4937. Given the different artifact signal response for participants S1, S6, and S7, we might expect diminished results when generalizing to test across participants.

To see how well the pattern of encoding regions observed in the single-participant experiment (Section 3.1) replicates to a new dataset, we performed the same encoding projection procedure in the NSD sample. Supplementary Figure 3 shows these results for all participants except S6, for which no voxels survived correction at FDR < 0.05. For participants S3, S5 and S7, we used more conservative thresholds of FDR < 0.0001, 0.005, and 0.001, respectively. To visualize the common brain regions across participants, we performed a one-sample, two-sided t-test using the average encoding weight maps for each participant. However, no voxels survived after correcting for multiple comparisons at a fairly liberal threshold, FDR < 0.05. Instead, we generated probability maps from the individual participant t-tests that display the probability of a voxel surviving correction across all participants. Given the large percentage of voxels that are significant for at least one participant, 57%, Figure 4C shows only the voxels with a positive probability of 0.25 or greater, and voxels with a negative probability of 0.125 or greater. Brain regions along the bilateral medial wall and in the posterior cingulate and superior parietal areas were positively correlated with heart period prediction for at least two participants. Bilateral temporal pole, anterior cingulate, superior frontal and left cerebellum brain regions were negatively correlated with heart period prediction for at least one participant. Thus we were largely able to replicate the cortical regions that regulate variation in instantaneous heart period. The variability in single trial encoding models, with strict corrections for multiple comparisons, is not surprising given the sample size used here, particularly since the goals of this study are prediction, not representational mapping.

#### 3.2.3 Generalization across participants

In order to see how well the decoding models work across participants, we tested generalization across the NSD participants, using the average trained model from one participant and testing it on every other participant. This allowed us to gauge the level of individual differences between decoding weight maps during the drive signal at +7.99 seconds. Figure 5A shows the average correlation between observed and predicted heart period for lag time shift +7.99 seconds for each training and testing participant combination of the group analysis. Darker colors represent lower prediction accuracy and lighter colors represent higher drive signal prediction accuracy. As anticipated, we are able to predict instantaneous heart period within participants more accurately than across participants, shown in the lighter colors along the diagonal and darker colors off the diagonal of Figure 5A. There are some exceptions for particular participants however. Decoding weight maps for S8 predicted instantaneous heart period for S5 (mean r = 0.1592) more accurately than their own heart period signal (mean r = 0.1215). Similarly, S1 predicted S7 more accurately (mean r = 0.1037 compared to mean r = 0.0572). The horizontal and vertical white lines separate the participants with expected artifact signals (S2, S3, S4, S5, S8) from those that behave somewhat as outliers (S1, S6, S7), with either low or no peak in prediction accuracy during the artifact time window. Indeed as expected, when the decoding weight maps from these outlier participants are used as the training model, drive signal prediction accuracy performs more poorly in general, as shown in the darker colors of the lower left section of the heat map. Comparatively, it is interesting to note that when the outlier participants are used as test participants (upper right rectangle), model performance is generally higher. Focusing on the participant combinations in the top left rectangle, we show that generalization across participants is somewhat possible, albeit with lower performance. However, it is also clear that certain participant combinations have better model performance, which emphasizes the individual nature of the decoding weight maps that predict instantaneous heart period.

**Figure 5:**
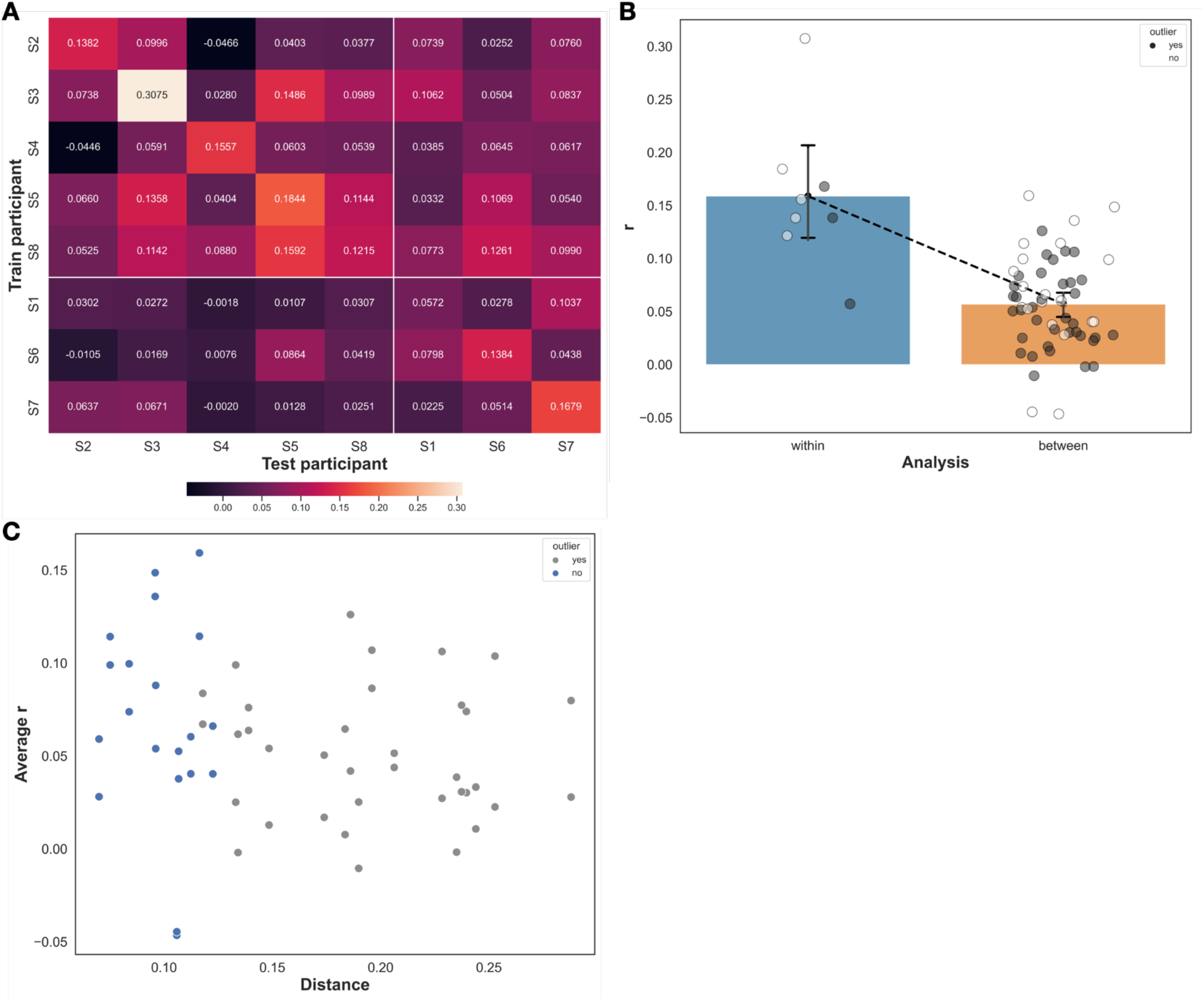
A) Heat map of mean out-of-sample Pearson correlation coefficient values from the group analysis across participants at lag time shift +7.99 seconds, with training participants on the y axis and testing participants along the x axis. B) Breakdown of within participant model r values (heat map diagonals) and across participant model r values (heat map off diagonals). Bars represent the mean r values, error bars show the 95% confidence intervals. Gray points indicate the presence of S1, S6 or S7 in the train-test participant combination. C) Scatter plot of interparticipant euclidean distance, on the x axis and the average decoding accuracy (r values) on the y axis for between participant models.

To get a clearer picture of this between-participant generalization, we plotted the within-and between-participant model accuracies separately in Figure 5B. Specifically, this compares the diagonal entries of the heat map (within-participant analysis) with the off-diagonal entries of the heat map (between-participant analysis), with gray dots representing a train-test participant combination that includes at least one outlier participant (S1, S6, S7). The within-participant analyses have higher mean prediction accuracies (mean r = 0.1588) compared to the between-participant analysis (mean r = 0.0570). The outlier participant combinations have slightly lower mean prediction accuracies on average, though there is not a dramatic difference.

One factor that might explain this variability in between-participant prediction accuracy is the overall similarity in their decoding weight maps. In other words, participants with more similar learned decoding models should also generalize better than pairs of participants with less similar models. To test this we calculated the Euclidean distance between decoding weight maps for each pair of participants and then saw how well this distance was associated with the ability of one participant’s model to predict the other’s. As a way of visualizing the association between the similarity of different participant weight maps and the average prediction accuracy, we generated the scatterplot shown in Figure 5C. Each point represents a between-participant pair (e.g. S1-S2), with gray points again representing participant combinations that contain one or more outlier participants. While there is no significant relationship overall (Pearson r = −0.2022, 95% confidence interval [−0.4416, 0.0641]), it appears that the weight maps of participants that are not outliers are more similar than outlier participants, regardless of prediction accuracy. Thus, similarity in maps may correlate with the ability to generalize, but the current sample may be too small to discern a reliable statistical effect

### 3.3 Across scanner generalization

Finally, we set out to see how well decoding maps generated from one MRI scanner, in this case the 7T scanner, generalizes to data from another scanner (the 3T, single-participant dataset). We did this by using the average NSD trained model (across all participants) and testing on the Human QA dataset runs for the drive signal timepoint (+7.99 seconds for NSD, +7.5 seconds for Human QA). This is a true generalization in that scanner strength, scan acquisition parameters and physiological recording methods were different across the two datasets. The correlation between observed and predicted heart period across runs ranged from r = −0.0568 to r = 0.4040, with a mean value of r = 0.1424 and 95% confidence interval [0.1058, 0.1775] (generated from 1000 bootstrap iterations), as shown in Figure 6. This demonstrates that our model is generalizable, though prediction accuracy is not as robust as in individual participant analysis.

**Figure 6:**
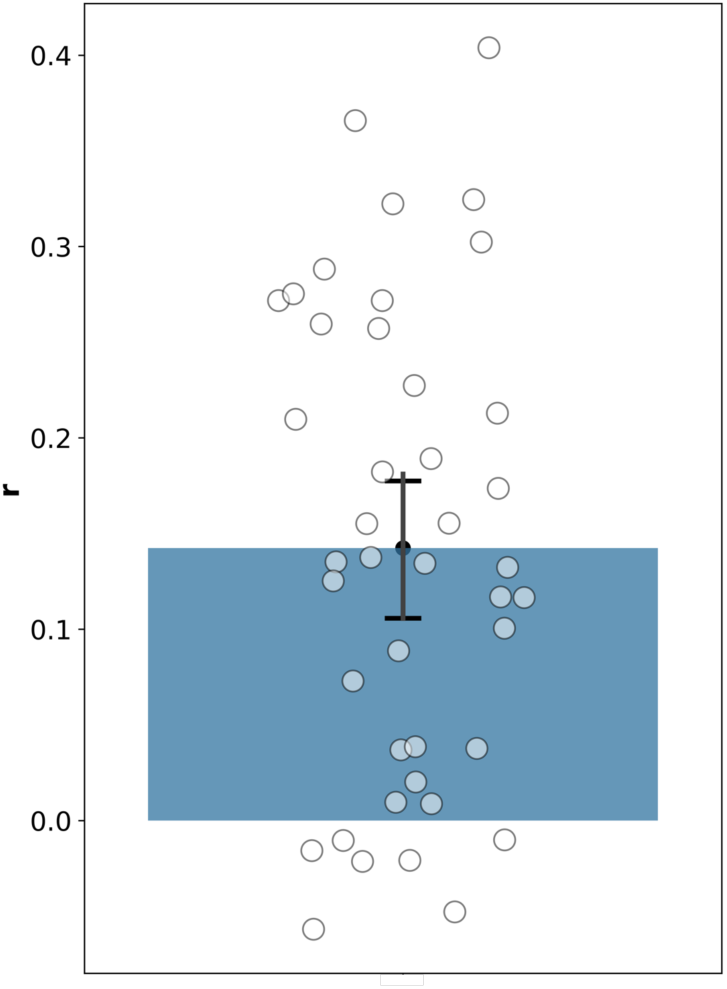
Correlation coefficient for each run from across scanner generalization models. The bar shows the mean r value, the error bars show the 95% confidence interval.

## 4. Discussion

Being able to reliably estimate the neural control of cardiac function, within individuals, would present a first step in developing clinical biomarkers of brain contributions to CVD. Using machine learning approaches optimized for high-dimensional datasets, we successfully predicted instantaneous heart period from whole brain hemodynamic data both within and across participants. We demonstrated these findings in two separate, highly sampled datasets that used different MRI and heart rate acquisition methods. We first observed robust prediction accuracy of instantaneous heart period for single participants. The strength of this within-participant effect has to do with the high statistical power of predicting events across individual BOLD samples, as opposed to the trialwise or blockwise event-related designs of typical fMRI experiments. This allows for reliable prediction of individual participant effects. We have shown that only a handful of runs are needed for reliably predicting instantaneous heart period. Models are also modestly generalizable across individuals, though with marginally lower prediction accuracies than within-participant models. Finally, brain regions in the parietal, frontal and temporal poles and in the anterior cingulate appear to reliably contribute to heart period prediction, suggesting that a network of cortical and subcortical brain regions work together to modulate the downstream brainstem’s control of heart period. Therefore, we have shown that dynamic fluctuations in the hemodynamic activity of upstream brain regions, comprising critical nodes of the visceral control circuit, track with and predict transient fluctuations of heart period.

Despite the individual differences in brain regions associated with heart period prediction across participants in our study, taken together, the overlapping regions from both datasets are largely in line with existing literature. Eisenbarth et al. (2016) demonstrated a multivariate pattern of social threat evoked fMRI activity that predicts heart rate [21]. The positive predictive weights for this model were located in the dorsal anterior cingulate and negative predictive weights in the medial prefrontal cortex, which coincide with our results from both datasets. In addition, Gianaros et al. (2004) studied the associations between heart period and regional cerebral blood flow (rCBF) during a working memory task. Negative correlations between heart period and rCBF in the insula, anterior cingulate also align with results from both of our datasets [35]. Similarly, Porro and colleagues (2003) also reported correlations between heart rate and fMRI activity during pain anticipation [36]. Brain regions in the parietal cortex were positively correlated while regions in the medial prefrontal cortex and cingulate cortex were negatively correlated, which again overlap with our findings. Finally, Critchley et al. (2000) found associations with heart rate and rCBF during motor and arithmetic tasks [37]. Their negative correlations in the medial prefrontal and cingulate cortices with heart rate also coincide with our results. Altogether, these collective findings emphasize the relationship between specific cortical and subcortical brain regions that might regulate the chronotropic aspect of cardiac activity, and by extension cardiovascular risk. Where our results extend this prior work is in showing that not only are these critical brain regions in higher-level cortex (i.e., upstream from the brainstem) associated with cardiac function, but can reliably predict it on a moment-by-moment basis and at the single-participant level.

With this in mind, there are two main methodological limitations to consider when interpreting the results of the present study. First, there are temporal differences between the parasympathetic and sympathetic heart rate responses [38] that we did not explicitly address in our analysis. It is possible that we only captured the parasympathetic response that occurs more immediately in response to stress. Intentionally incorporating these two autonomic heart rate responses into our model and investigating how this may alter both the accuracy of prediction and the brain regions involved would be a worthwhile future endeavor. Second, we treated task and rs-fMRI data identically, as the tasks from both datasets were not explicitly designed to evoke changes in heart rate, but any spurious changes in cardiac activity only increased usable variance for our models to pick up on. It is still possible, however, that there are different brain processes across task-states that may have had some influence on our models. Follow-up work will explore this difference between situations where there may be more top-down control of cardiac function (e.g., tasks) from those where the system may be in a more passive state (e.g., rest).

Also, while our model predictions showed fairly high consistency at predicting heart period across participants, it is worth noting that the encoding maps showed substantial variability as well. In some ways this is not surprising. The decoding model used here was optimized for prediction, not localization, and uses correlated patterns across the entire brain to build its prediction. This can introduce substantial variability in the decoding weights, and consequently, the encoding projections used to localize regions of influence. Our results in the 7T dataset suggest that any questions on precise localization of control should rely on much larger datasets where consistent patterns across individuals can be more reliably discerned.

Finally, it is important to keep in mind that our sample consisted of only healthy, young to midlife individuals. While the cardiovascular health of our sample was not directly tested, it is very likely that any cardiovascular disease risk in this sample is minimal. Thus, if we hope to translate the current approach to a meaningful measure of potential cardiovascular disease risk, follow up work should investigate whether cortical control of heart period that we observe here persists in healthy older populations that often have higher risk of cardiovascular disease, as well as in populations with overt cardiovascular disease or other comorbidities. In order to make these translational steps, we must first prove that it is possible to measure the neural regulators of heart rate. The current study provides this proof of concept that can be leveraged by future studies.

Despite these limitations, our study makes clear the feasibility of measuring the cortical and subcortical control signals, including brainstem areas, that are associated with variation in cardiac function in humans. To expand on the relevant immediate next steps, repeating our analysis with extended lag time shifts to look at predicting the sympathetic response and trying to tease out the distinction between the two responses could provide some additional nuances to our current results. It would also be valuable to examine any differences between the brain regions most important for parasympathetic versus sympathetic heart rate responses. Performing a simulated lesion analysis, by removing certain brain regions in turn, would be another method of evaluating the relative importance of individual brain regions (or networks) contributions. Finally, replicating this analysis on a much larger datasets, of hundreds or thousands of participants, would boost our ability to reliably localize consistent regions of control in the normative human brain. All of these approaches reflect important next steps in our work.

## 5. Conclusion

We have added to the growing existing body of literature looking at brain-heart connections, showing that heart period is reliably predicted by brain activity, even at the single-participant level, within cortical and subcortical regions that overlap with visceral control circuit networks. These results offer a step towards the development of a clinically applicable brain-based biomarker for CVD risk. Ultimately, continued work to elucidate the role of the brain’s cortical control mechanisms on cardiovascular function and disease risk is essential to further the development of novel therapies and prevention strategies for CVD.

