## Supplementary Figure 1 for "Cortical and subcortical brain networks predict prevailing heart rate"

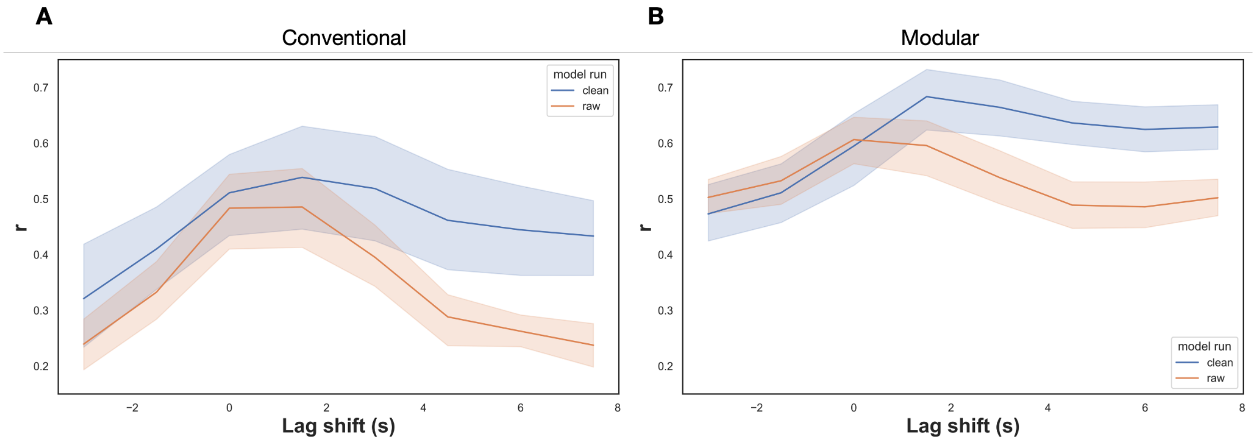


**Supplementary Figure 1:** Mean out-of-sample Pearson correlation coefficient of predicted and observed instantaneous heart rate across four sessions (12 runs total) for each lag time shift for both the clean fMRI signal and the artifact fMRI signal for the conventional analysis (A) and modular analysis (B) approaches. Shaded regions represent 95% confidence intervals (calculated using 1000 bootstrap iterations).
