## Supplementary Figure 2 for "Cortical and subcortical brain networks predict prevailing heart rate"

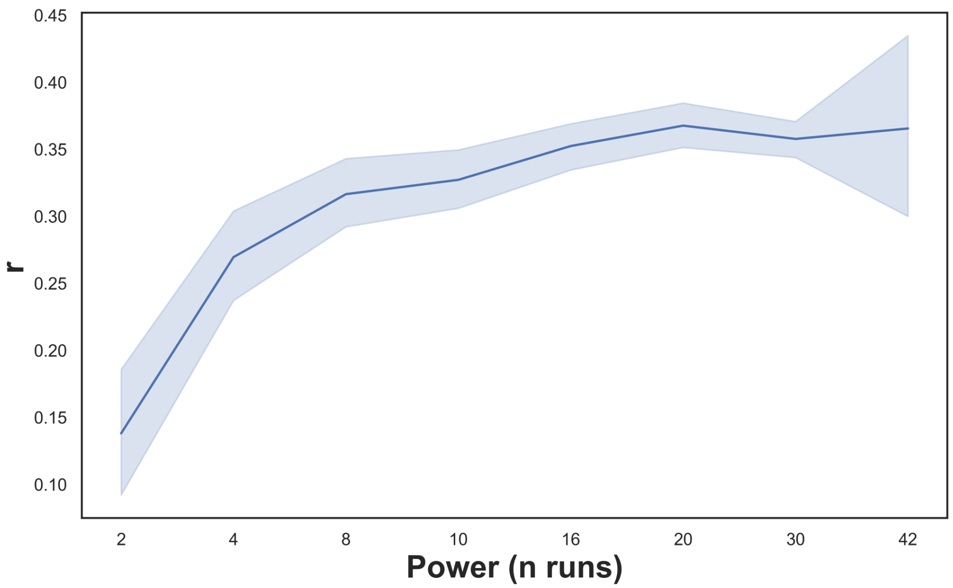


**Supplementary Figure 2:** Mean out-of-sample Pearson correlation coefficient of predicted and observed instantaneous heart period across different sample sizes of Human QA dataset runs. For sample sizes n = 2 through n = 30, results are averaged across 40 iterations with randomly selected runs. The shaded region represents 95% confidence intervals (calculated using 1000 bootstrap iterations).
