## Supplementary Figure 3 for "Cortical and subcortical brain networks predict prevailing heart rate"

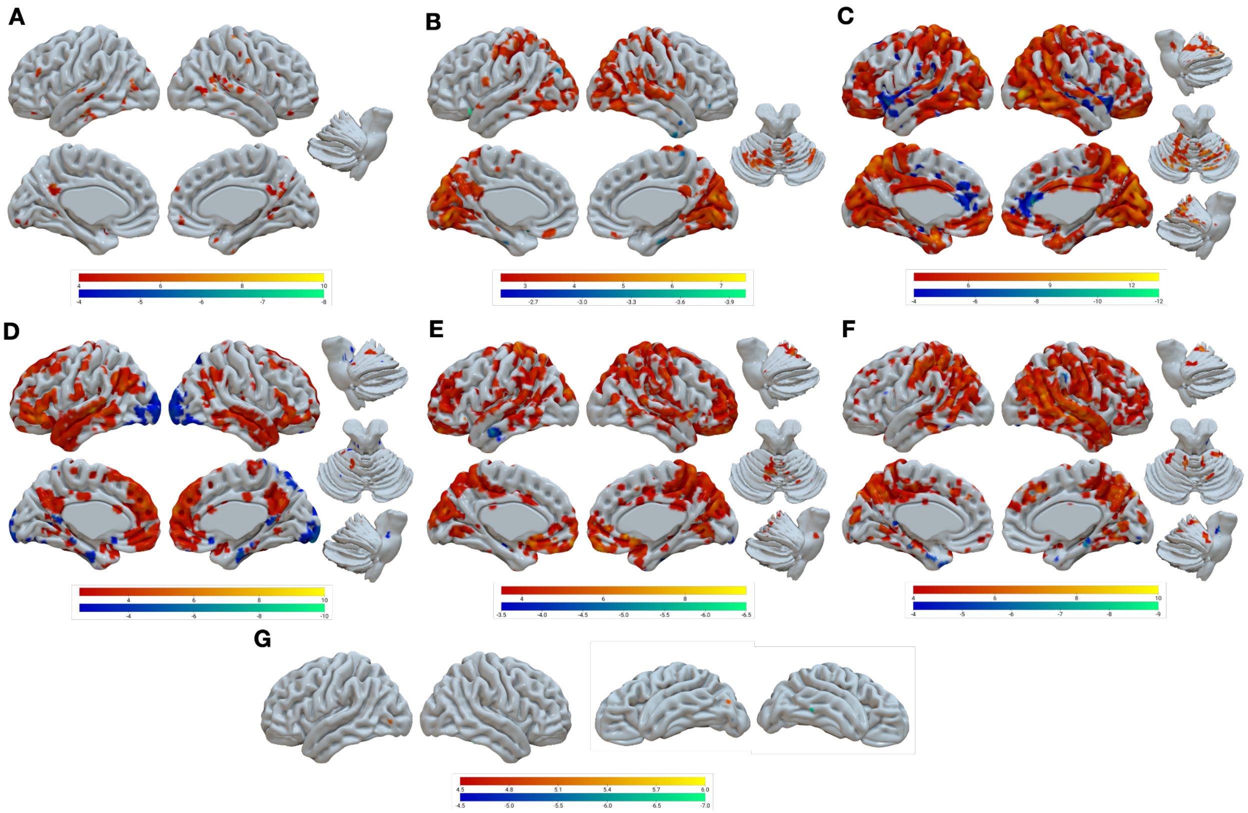


**Supplementary Figure 3:** One-sample, two-sided t-tests of encoding weight maps of instantaneous heart period prediction for each NSD participant at lag time shift +7.99 seconds. A) S1, FDR < 0.05, B) S2, FDR < 0.05, C) S3, FDR < 0.0001, D) S4, FDR < 0.05, E) S5, FDR < 0.005, F) S7 FDR < 0.001, G) S8, FDR < 0.05. Note: S6 is not included since no voxels survive correction at FDR < 0.05. Positive weights are shown in red-yellow. Negative weights are shown in blue-green.
