## Supplementary Materials for "Cortical and subcortical brain networks predict prevailing heart rate"

Prediction of instantaneous heart period:

We first set out to show that the modular approach, where we average decoding maps across runs to predict instantaneous HR in a hold out run (see Section 2.1.4), performs as well as a more conventional approach where a single decoding map is generated from data aggregated across training set runs. This was done on a subset of the single-participant dataset (four sessions, 12 runs total). Supplementary Figure 1 shows the Pearson correlation coefficient values for both analysis approaches. The modular approach achieves qualitatively similar results, if not has overall higher prediction accuracy, than the conventional modeling approach. We found this to be acceptable to move forward with the modular approach for all subsequent analyses.
